# Real-time, *in situ* fluorescence and optical density measurements of liquid cultures in simulated microgravity

**DOI:** 10.64898/2026.03.23.713711

**Authors:** Stephen Lantin, Mayur Bansal, Hal S. Alper, Jessica Audrey Lee

## Abstract

As human space exploration expands to the Moon, Mars, and beyond, there is a growing need to study the effects of altered gravity on the microbial systems that we will bring with us for life support. Because spaceflight experiment opportunities are rare and resource-intensive, most space biology experiments are conducted using ground-based simulators. The most common microgravity simulator for microbial experiments, the rotating wall vessel, can approximate the low-shear and low-turbulence conditions that characterize microgravity. However, current designs do not allow for real-time measurement of growth or metabolic activity during rotation: experiments require destructive sampling or disruption of the microgravity simulation conditions. Here, we describe the development of an *in situ* spectroscopy system compatible with the Cell Spinpod rotating wall vessel, which enables measurement of both optical absorbance and fluorescence with high temporal resolution, producing growth curves similar to those from an off-the-shelf plate reader. These results are validated using two common microbial hosts: *Escherichia coli* and *Saccharomyces cerevisiae*. The Spinpod Optical System has the potential to diversify the types of microbiology experiments possible in simulated microgravity, allowing the measurement of not only growth curve parameters but also metabolic activity, gene expression, or community dynamics. It thus has the potential to improve the quality of experiments seeking to characterize microbial responses to spaceflight conditions.

## 1. Introduction

As human presence in space expands through long-duration missions, understanding how single-celled organisms respond to space-associated stressors is increasingly vital. Microbes will be essential to sustained human habitation of the Moon, Mars, and deep space through their roles in bioregenerative life support systems, including waste processing, oxygen generation, and the in-situ production of food, nutrients, and medicine [1–4]. Their ability to function reliably in altered gravity will therefore directly impact the success of future missions. Investigating how these organisms respond to the stress of altered gravity provides both practical benefits for space habitation and deeper insight into the fundamental interactions of organisms with their physical environments. However, there are few physical systems that allow for terrestrial interrogation of these effects in real time, while organisms are experiencing simulated microgravity.

Environmental conditions associated with spaceflight induce physiological changes at all levels of biocomplexity. Reduced gravitational forces result in a decrease in both density-driven convection and particle sedimentation, causing diffusion and surface tension effects to dominate transport processes. Microbes in liquid culture therefore experience very low shear stress and reduced fluid mixing, which can alter local chemical concentration gradients around cells [5–10]. Microbial behavior has been observed to change as a result of both spaceflight and ground-based microgravity simulation conditions [11–13], suggesting increased antibiotic resistance and virulence [9,14–18], biofilm formation [19,20], horizontal gene transfer [21], secondary metabolite production [22–24], altered gene expression profiles [9,16,25–27], and altered growth parameters such as lag time, growth rate, and final biomass [24,28,29]. While research has led to key fundamental understandings about microgravity-sensitive responses in some model organisms, there is still uncertainty regarding the correspondence between ground-based and spaceflight investigations [30,31]. In addition, much work remains to be conducted on the diversity of other microbial species and multispecies co-cultures. The inaccessibility of appropriate experimental hardware for spaceflight investigations is a well-known barrier to conducting rigorous research and allowing new researchers to contribute to the field.

### 1.1 Challenges

Microgravity cannot be fully reproduced in any lab on Earth’s surface. To simulate microgravity in experiments aimed at understanding organisms’ response to altered gravity, the microgravity simulation method must model the physical aspects of the microgravity environment to which the organisms respond. Microorganisms such as bacteria and yeast are believed to be unable to sense the force of gravity directly [24,30]; microgravity simulation for microbes focuses instead on reproducing the quiescent, non-mixing, low-shear fluid environment of microgravity, and removing the ability of cells to settle through the fluid [8,24,32–38].

This simulated microgravity environment has most commonly been generated using Rotating Wall Vessel (RWV) bioreactors: cylindrical culture vessels filled with liquid culture medium (no gaseous headspace) are rotated during the experiment. The High Aspect Rotating-Wall Vessel (HARV) developed at NASA was the first of these designs, featuring a horizontal axis of rotation and the lack of internal stirring vanes [39,40]. By taking into account several parameters, including differences in density between cells and fluid, fluid density, and fluid viscosity, cell terminal velocities may be achieved in an RWV such that shear and turbulence are minimized. HARV-type RWVs have been among the most commonly used by microgravity researchers in past decades [8]. To improve gas exchange for aerobic cultures, alternative designs with coaxial tubular oxygenator were developed [41,42]; shear is likewise minimized if the inner oxygenator tube is rotated at the same angular speed as the outer vessel. Several additional models of rotational culture systems have also been developed, including the use of the long, relatively small-diameter Fluid Processing Apparatus as an RWV, and a relatively new, optically clear commercial product called the Cell Spinpod. Other instruments have been developed to simulate microgravity in ground-based studies, including 3D clinostats and the random positioning machine [43,44]. While these are more effective at averaging the gravity vector in all directions, they can produce turbulent flows in the fluid culture medium, which makes them inappropriate for experimentation seeking to investigate the effect of the quiescent microgravity environment on microbial cells.

An important limitation of existing rotating wall vessel designs is the difficulty of monitoring the activity of organisms during rotation. Experimenters traditionally manage this challenge by stopping rotation mid-experiment to remove samples, by initiating many replicate cultures at once and harvesting a few at each timepoint (destructive sampling), or by constraining analyses to endpoint measurements [26,45,46]. All of these approaches result in some loss of science: stopping rotation can result in altered fluid flows, compromising the fidelity of the microgravity simulation; destructive sampling is resource-intensive and can introduce additional experimental variability because time course measurements are not all made from the same sample; and dependence on endpoint measurements can miss important temporal dynamics during the course of growth, such as carrying capacity, intrinsic growth rate, doubling time, and initial population size [47].

Here we describe an easily reproducible, custom-built experimental system designed to overcome these limitations by measuring optical density and biological fluorescence *during* microbial growth in simulated microgravity. This system uses the Cell Spinpod, a commercially available single-use rotating wall vessel [48]. Two features of Cell Spinpods make it ideal for making real-time optical measurements during rotation: spinpods are optically clear; and a spinpod is seated onto two rotating bars rather than being mounted to a rotator via an attachment on its rear wall, allowing equal access to both the front and back of the spinpod for measurements of light transmission during rotation. Our custom system adds light sources, optical measurements by spectrometer, and data analysis and standardization. Our aim was to build a system that could measure both optical density and biological fluorescence well enough to compare growth and fluorescent protein expression curves in spinpods to similar curves measured in a standard benchtop multiwell plate reader.

### 1.2 Technical Requirements

For effective liquid culture microgravity experimentation on Earth, we require a system that satisfies four technical requirements (TRs):

[TR1] maintains biocompatibility for a variety of microbial cultures across full growth curves;

[TR2] reproduces microgravity-like conditions;

[TR3] acquires reliable, real-time fluorescence emission and optical density measurements;

[TR4] does not disrupt the simulated microgravity conditions during measurement.

## 2. Materials & Methods

### 2.1 Hardware

To contain liquid cultures for experimentation, the described setup uses Cell Spinpods (Cell Spinpod, LLC, Chapel Hill, NC, USA), which are optically clear, keep liquid microbial cultures contained, and are designed for rotational culture. Spinpod vessel walls are composed of polystyrene and polycarbonate, materials whose biocompatibility and nonreactivity with a range of liquid media are known [48–50]. If the selected medium is moderately viscous, growth substrates will diffuse as they are locally depleted to permit cell growth. Therefore, [TR1] is satisfied. Spinpods filled with microbial cultures were set in Carrier Extenders (Cell Spinpod, LLC) and placed on the rollers of a rotator (Mini-Rotator, Cell Spinpod, LLC; **Fig. 1**). (The system can also be adapted to the larger Thermo Scientific Bottle/Tube Roller.) Carrier Extenders provided a way of mounting the light source near the sample for optical density measurements (see below). Because the spinpod setup tended to migrate backward along the rollers during rotation in a way that affected the distance between the sample and the optical probe, several empty Carrier Extenders were placed on the rollers behind the spinpod and light source setup, as spacers, to prevent backward migration. To accommodate the Carrier Extenders, the roller arm of the Mini-Rotator (typically used to keep spinpods in place) was removed. For experiments shown in this manuscript, rollers were made to rotate with a setting of 25 RPM (resulting in a rotation rate for the spinpod itself of 20 RPM, due to the difference in diameter between the rollers and the spinpod with casing). The spinpod and rotator were maintained in a darkened incubator to eliminate background light and maintain temperature; power and optical cables were fed to the outside of the incubator, where the spectrometer, a laptop computer, and PHOS-4® light source were maintained. A bill of materials is provided in *Supplemental File 1*; CAD drawings are available in a repository at OSF.io [51] (doi: 10.17605/osf.io/b49gm).

**Figure 1.**
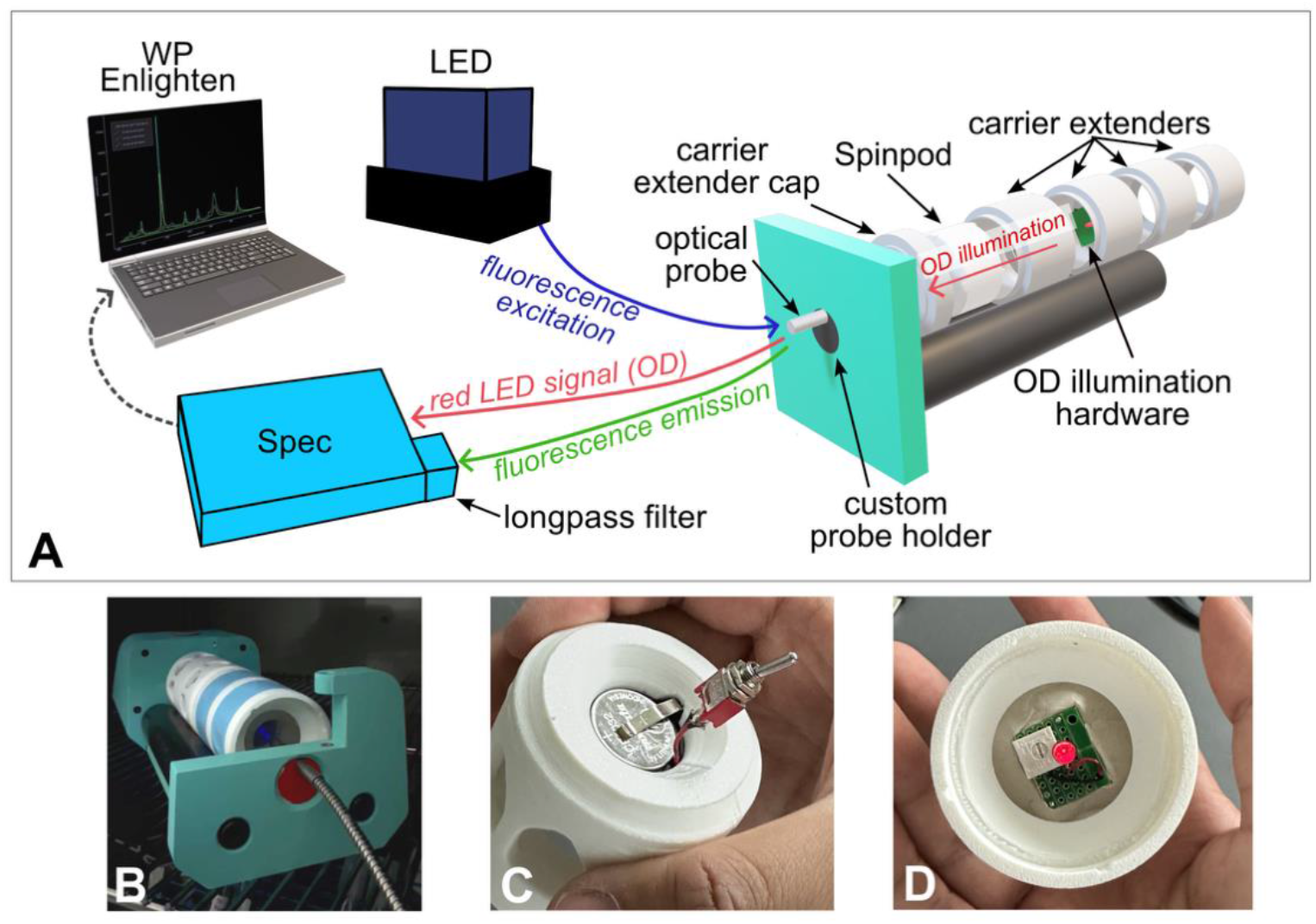
A) A close-up, exploded view of the spinpod optical system and diagram of operations. To take fluorescence measurements, excitation light first passes through a bidirectional optical probe from the LED light source to illuminate liquid cultures encased in a spinpod. Fluorescence emission then returns through the optical probe to the spectrometer for measurement, through a longpass filter that reduces signal from the excitation wavelength. To take optical density measurements, a red LED mounted toward the optical probe but on the opposite side of the spinpod illuminates the spinpod (“OD illumination hardware”). This light passes through the spinpod and to the spectrometer without filtering. The spinpod is housed in two joined Carrier Extenders with additional Carrier Extenders placed inline as spacers. B) Photograph of the system diagrammed in panel A; optical probe is in front. Tape has been applied to the outside of carrier extenders to shield the optical path from external light. C) and D) Rear and front close-up views, respectively, of the OD illumination hardware.

#### 2.1.1 Optical density components

To provide suitable illumination for the optical density measurement, a red LED was mounted facing the probe on the opposite side of the spinpod in a Carrier Cap (Cell Spinpod, LLC). Mounting putty (Gorilla Brand, product 168) was used to mount the perf board to the back of the Carrier Cap, with a CR2032 coin battery and holder fitted into the viewport; note that a small portion of the Carrier Cap needed to be whittled away with a knife or other sharp tool for best fit (**Fig. 1C**). A red indicator LED with peak emission at 695 nm was chosen in order to avoid conflict with the wavelengths involved in excitation and emission of yellow and green fluorescent markers. The intensity of the LED was low enough that a 10-second integration time (necessary for detecting fluorescence) could be used for data acquisition, without saturating the detector. A power switch allowed the light to be turned on and off manually. A schematic and bill of materials can be found in the Supplemental Data.

#### 2.1.2 Fluorescence optical components

Fluorescence measurements used a precision LED source (PHOS-4^®^, In Tandem Designs Pty Ltd, Warrnambool, VIC, Australia), with light transmitted by the fiber optic cable (*Section 2.1.3*). For all experiments shown here, only the blue light (P4-VIS-475-BBL); *λ*_peak_ = 475 nm) was used, at maximum intensity. A 487 nm longpass filter (reducing light with wavelengths shorter than 487 nm) was included in the optical path between sample and spectrophotometer, in an attached filter holder/cuvette holder unit, to reduce any light from the excitation source that was reflected back by the sample.

#### 2.1.3 Signal transmission

To transmit the optical signal, a Y-shaped fiber optic bundle reflection probe was used (Ø600 μm, 0.39 numerical aperture, single-fiber spectrometer leg and fiber bundle light source leg), with a Ø1/4” probe on the sample leg and SMA connectors on the light source leg and spectrometer leg. The probe was fixed normal to the center of the spinpod’s viewport. A custom probe holder was designed and fabricated by 3D printing and inserted in the rear porthole of the Mini-Rotator to keep the optical probe in place (**Fig. 1**; Design files included in Supplemental Information). The detector end of the cable fed into a visible-range spectrometer (WP VIS, Wasatch Photonics, Logan, UT, USA).

### 2.2 Strains and Media

A modified *E. coli* BL21 (DE3) strain was used for single-species bacterial experiments measuring optical density. A codon-optimized mCherry gene and a kanamycin resistance gene were inserted into the *recA* locus. Additionally, this strain also carried one plasmid: CDuet™-1 vector carrying chloramphenicol resistance. (Fluorescence was not measured in this strain.) Cultures were inoculated from a colony into Luria-Bertani (LB) broth with 50 µg/mL kanamycin and 25 µg/mL chloramphenicol and grown overnight, shaking at 200 RPM, at 37 °C. They were then passaged at a 1:100 dilution into supplemented M9 minimal medium. This consisted of 1 x M9 mineral salts (final concentrations: 12.8 g/L NaHPO_4_ ⋅ 7 H_2_O, 3 g/L KH_2_PO_4_, 0.5 g/L NaCl, 1.0 g/L NH_4_Cl), supplemented with 1 µM MgSO_4_, 0.1 µM CaCl_2_, 1% casamino acids, 20 g/L glucose, 50 µg/mL kanamycin, and 25 µg/mL chloramphenicol, and sterilized by filtration through a 0.2 µm filter. The culture in M9 was again grown overnight, shaking at 200 RPM at 37 °C, to reach stationary phase. To initiate a spinpod experiment, this stationary-phase culture was then diluted in fresh supplemented M9 minimal medium to reach a starting OD_600_ (as measured in a benchtop spectrophotometer) of 0.025 in a final volume of 5 mL, and this dilute culture was immediately filled into a spinpod through the fill port using a syringe and needles, mounted into the optical measurement system, and rotated.

The *S. cerevisiae* strain was based on BY4741 and contained genomic integrations of *mjDOD* and *CYP76AD* at the URA3 locus for betaxanthin production. An additional plasmid based on the pRS vector backbone provided geneticin resistance. Experiments were initiated by inoculating colonies from a plate into liquid Yeast Peptone Dextrose medium (YPD; 10 g/L yeast extract, 10 g/L peptone, 20 g/L glucose) supplemented with 200 µg/mL geneticin, and growing overnight, shaking at 200 RPM, at 30 °C. Cells were then transferred to yeast Synthetic Complete (SC) medium with Complete Supplement Mixture (CSM; adenine 10mg/L, L-arginine HCl 50mg/L, L-aspartic acid 80mg/L, L-histidine HCl 20mg/L, L-isoleucine 50mg/L, L-leucine 100mg/L, L-lysine HCl 50mg/L, L-methionine 20mg/L, L-phenylalanine 50mg/L, L-threonine 100mg/L, L-tryptophan 50mg/L, uracil 20mg/L, L-tyrosine 50mg/L, L-valine 140mg/L, total 790 mg/L) with 6.7 g/L yeast nitrogen base, 20 g/L glucose, and 200 µg/mL geneticin. As with *E. coli* experiments, spinpod experiments with yeast were initiated by diluting stationary-phase culture grown in supplemented SC into fresh supplemented SC to reach a starting OD_600_ (as measured in a benchtop spectrometer) of 0.025 in a final volume of 5 mL. The freshly diluted culture was immediately filled into a spinpod, mounted into the optical measurement system, and rotated.

### 2.3 Growth experiment setup

All measurements were made using WP ENLIGHTEN™ software (Wasatch Photonics, Logan, UT, USA), via laptop connected to the spinpod optical setup (**Fig. 1**). Optical acquisition settings were as follows: integration time 10 seconds; boxcar smoothing 4 pixels; no scan averaging. Growth experiments in the spinpod optical system were initiated by inserting the freshly inoculated spinpod into the carrier extenders and placing on the rotator with both light sources off. A “dark” reading was then taken and stored. Then the light sources were turned on as appropriate (red only for OD-only experiments; both red and blue for OD + fluorescence experiments). Both light sources were kept on at constant intensity for the full duration of the experiment. Batch acquisition of spectra was initiated, taking 3 measurements in quick succession at each timepoint, with one timepoint every 10 minutes. During the experiment, the incubator was kept closed to eliminate background light and set at the optimal temperature for the organisms in the experiment (37 °C for *E. coli*, 30 °C for *S. cerevisiae*). All growth experiments were conducted for 24 hours.

### 2.4 Standard curves

Standard curves were generated by making diluted suspensions of cells and measuring the same samples in both the spinpod optical system and in a standard benchtop spectrophotometer (Genesys 10S UV-VIS, Thermo Scientific). For *E. coli*, stationary-phase cell cultures grown in supplemented M9 medium were diluted in fresh supplemented M9 to concentrations of 1.0, 0.8, 0.6, 0.4, 0.2, and 0.1 of the stationary-phase cell density. Each dilution was filled into a spinpod, the spinpod mounted in the spinpod optical system, and 4 replicate measurements were taken of each dilution as well as sterile M9 without cells, using the same optical settings as for growth experiments (see *Section 2.3*). The same samples were also aliquoted into cuvettes (1 cm path length) and OD_600_ was read on the spectrophotometer. All measurements were made within 20 minutes of dilution to avoid complications due to microbial growth in the fresh medium. For *S. cerevisiae*, stationary-phase cell cultures grown in supplemented SC medium were diluted in phosphate-buffered saline (PBS) solution (pH 7.1) to concentrations of 1.0, 0.9, 0.8, 0.7, 0.6, 0.5, 0.4, 0.3, 0.2, and 0.1 of the stationary-phase cell density. Each dilution was filled into a spinpod and into a cuvette and measurements made as for *E. coli*. No growth was expected during the course of measurement because the PBS contained no carbon.

## 3. Theory/calculations

To acquire fluorescence measurements, a PHOS-4^®^ LED light source emits blue light into one end of a bifurcated fiber optic cable, which is connected via optical probe to a rolling spinpod. Liquid cultures contained in the spinpod fluoresce at a longer wavelength back into the optical probe, through the other bifurcation of the fiber optic cable, and into the Wasatch Photonics spectrometer (**Fig. 1A**). Fluorescence intensity can be visualized and quantified live via the WP ENLIGHTEN™ software. To acquire optical density measurements, the PHOS-4^®^ LED light source is turned off and light from a red LED passes through the spinpod, into the optical probe and fiber optic cable, and into the Wasatch Photonics spectrometer.

Collected optical data were analyzed using a set of custom R scripts; scripts and associated input files from both the standard curves and the growth curves are available in a repository at OSF.io[51] (doi: 10.17605/osf.io/b49gm). In brief, each spectrum from each timepoint was read into R as a data frame, and for each timepoint the heights of the peaks at the wavelengths of interest were extracted. For optical density, the peak of interest was 695.08 nm. For betaxanthin fluorescence, the peak of interest was 510.14 nm. In addition, a “baseline” peak (385.51 nm) was also extracted and tracked to monitor for potential drift in the detector or light source. For optical density, a standard curve was constructed for each strain to correlate ground-truth OD_600_ (as read in a benchtop spectrophotometer) to the optical density at 695 nm as measured through a spinpod. Because optical density is typically correlated to the negative logarithm (base 10) of absorbance, the following formula was used:

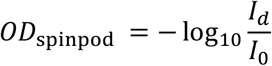

Where *OD*_spinpod_ = the optical density of the sample in the spinpod, *I*_*d*_ = the height in counts of the 695 nm peak measured through the sample of interest, and *I*_0_ = the height in counts of the 695 nm peak measured through a blank spinpod (medium only). For standard curves, *I*_0_ was measured in a blank sample; for growth curves, *I*_0_ was taken as the height of the 695 nm peak at the initial timepoint, while the density of cells was still below the limit of detection. The lm() function in R was used to calculate the correlation between *OD*_spinpod_ and benchtop OD_600_ by least-squares regression, using the data from the cell dilutions. This linear model was then used to calculate the “equivalent OD_600_” for all growth curves conducted in spinpods. Note that using 695 nm peak values in the calculation of *OD*_spinpod_ instead of 600 nm is reasonable as strains do not have significant pigment absorption in either wavelength; **Fig. 2B** demonstrates a strong linear correlation.

**Figure 2.**
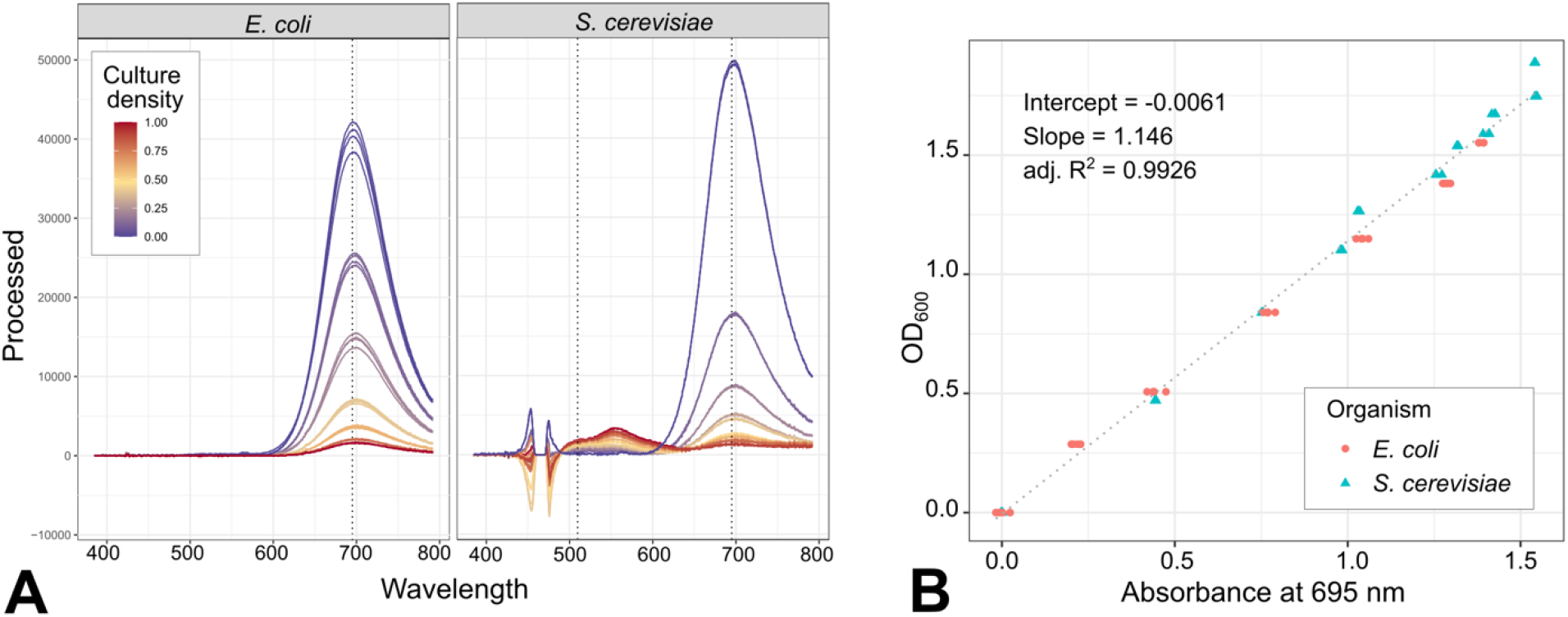
A) Raw optical spectra from *E. coli* and *S. cerevisiae* standard curves. Dotted lines at 695.08 nm indicate peak wavelength used for absorbance calculations; dotted line at 510 nm in *S. cerevisiae* subplot indicates betaxanthin fluorescence emission peak wavelength. Irregular peaks <490 nm in the *S. cerevisiae* plot are due to the subtraction of the peaks in the “blank” spinpod, which included some reflected light from the 475 nm excitation source. B) Correlation between optical density measured in the spinpod optical system at 695 nm (*x*-axis) and optical density measured in a spectrophotometer at 600 nm (*y*-axis); *E. coli* were measured in supplemented M9 medium and *S. cerevisiae* in phosphate-buffered saline (color and shape indicate organism type).

Statistical comparison of standard curves made with two different organisms was conducted by analysis of covariance (ANCOVA), using the aov() function in the “car” package in R. Before conducting ANCOVA, the assumption of independence of variables was tested using the aov() function to look for significance of a model explaining one variable by the other; and the assumption of homogeneity of variance was tested using the leveneTest() function in the “car” package. Standard curves were tested for whether they were statistically different from a *x* = *y* line using the linearHypothesis() function in the “car” package.

## 4. Results

### 4.1 Optical density measured in the spinpod system is linearly correlated to standard spectrophotometer optical density measurements

To benchmark the performance of the spinpod optical system relative to a standard benchtop spectrophotometer, we compared absorbance of *E. coli* and *S. cerevisiae* cultures at 695 nm, as measured in a spinpod (*OD*_spinpod_), to the optical density at 600 nm of the same sample measured in a cuvette in a commercial spectrophotometer (OD_600_). Calculating the correlation also allowed us to translate spinpod optical data into familiar units for use in growth curve analysis.

We found that the relationship between *OD*_spinpod_ and OD_600_ was linear for the full range of cell densities relevant to our experiment, and the correlation was strong (**Fig. 2**). While there was a small statistically significant difference between the standard curve generated with *S. cerevisiae* and that generated with *E. coli* (ANCOVA Pr(>F) = 0.00321), it was not possible to determine whether this was due to differences between organisms or simply variation between experiments conducted on different days. Calculated separately, the slope of the *E. coli* standard curve was 1.080424, intercept 0.022694, and adjusted R-squared 0.9973; the slope of the *S. cerevisiae* standard curve was 1.18227, intercept -0.02799, and adjusted R-squared 0.9927. Because of the similarities in values, we combined the datasets to generate a standard curve with a slope of 1.146, an intercept of 0.0061, and an adjusted R-squared of 0.9926 (**Fig. 2**). Although the slope was similar to 1, it was significantly different from 1, for both the combined standard curve and each of the species-specific standard curves (Pr(>F) < 2.2 × 10^-16^).

### 4.2 Optical density measurements in spinpods saturate at high cell densities similarly to spectrophotometer measurements

We also analyzed the correlation between *A*_spinpod_ and cell density. For this analysis, we expressed the cell density in each analyzed sample as a fraction of the total cell density in the original, undiluted, stationary-phase culture, since all analyzed samples were dilutions of the stationary-phase culture. Note that the maximum cell density (1.0) for *S. cerevisiae* was ∼8×10^7^ cells/mL and maximum cell density (1.0) for *E. coli* was ∼9×10^8^ cells/mL, but the *S. cerevisiae* culture had a higher OD_600_ than that of *E. coli*.

We found that *OD*_spinpod_ was not linearly correlated to cell density for the full standard curve: at high cell densities, *OD*_spinpod_ increases more slowly than cell density does (**Fig. 3**). This effect is not unique to *OD*_spinpod_; it is very similar to the relationship observed between OD_600_ and cell density (**Fig. 3**), and to the general non-linearity of optical density with cell density at high cell densities that has been documented frequently in the literature [52,53]. One way to express this is to examine how well the goodness of fit of the linear model, expressed as the adjusted R-squared value, changes as observations from higher cell densities are added to the model (**Fig. 3**). For both species, the linear relationship between *OD*_spinpod_ and cell density deteriorates with increasing cell density, whereas the relationship between *OD*_spinpod_ and OD_600_ remains strong regardless of cell density.

**Figure 3.**
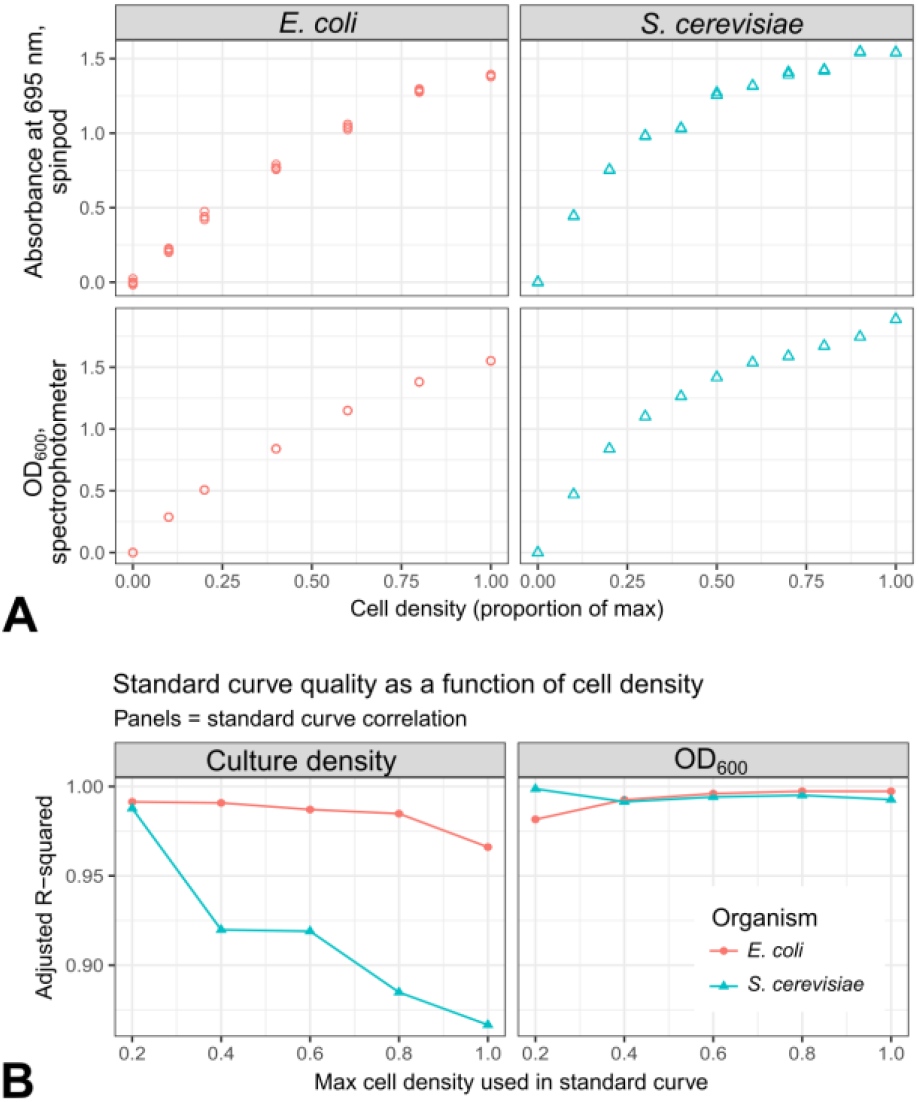
A. Relationship between *OD*_spinpod_ and cell density (top) and relationship between OD_600_ and cell density (bottom). B. Adjusted R-squared values for (left) *OD*_spinpod_∼cell density correlation and (right) the *OD*_spinpod_∼OD_600_ correlation, as a function of the cell density. The value on the x-axis denotes the highest cell density used in the correlation, i.e. an *x*-axis value of 0.8 means that all samples with cell density of 0.8 and below were used in the correlation.

### 4.3 Fluorescence measurements of constitutively expressed pigments are correlated with optical density in spinpods, at a given growth stage

To demonstrate the use of the spinpod optical system for measuring fluorescence, we used *S. cerevisiae* cells engineered for constitutive expression of betaxanthin, a yellow-orange pigment that fluoresces green (maximum emission wavelength: 510 nm) when excited with blue light [54]. We hypothesized that in the standard curve, in which cells came from a uniform population at the same growth stage (stationary phase), betaxanthin fluorescence would correlate to total cell density.

We found that in the spinpods, we were able to detect betaxanthin fluorescence as a peak observed in the spectrum between 500 and 600 nm in the *S. cerevisiae* samples that increased with increasing cell density (and confirmed that no such peak was observed in the *E. coli*, which did not express betaxanthin) (**Fig. 2**). In our system, the peak’s maximum value was closer to 550 nm than to the typically reported 510 nm, but we used the peak height at 510 nm for all analyses for agreement to prior experiments conducted with these strains. We found that the peak height at 510 nm correlated well with OD_600_ in the same samples (**Fig. 4**), indicating that fluorescence could be used as an alternative method of monitoring growth (assuming that growth conditions do not affect per-cell betaxanthin expression during the course of the experiment). We also found that peak height at 510 nm saturated at high cell densities, following a similar trend to that observed in OD_600_ and *OD*_spinpod_ (**Fig. 2**).

**Figure 4.**
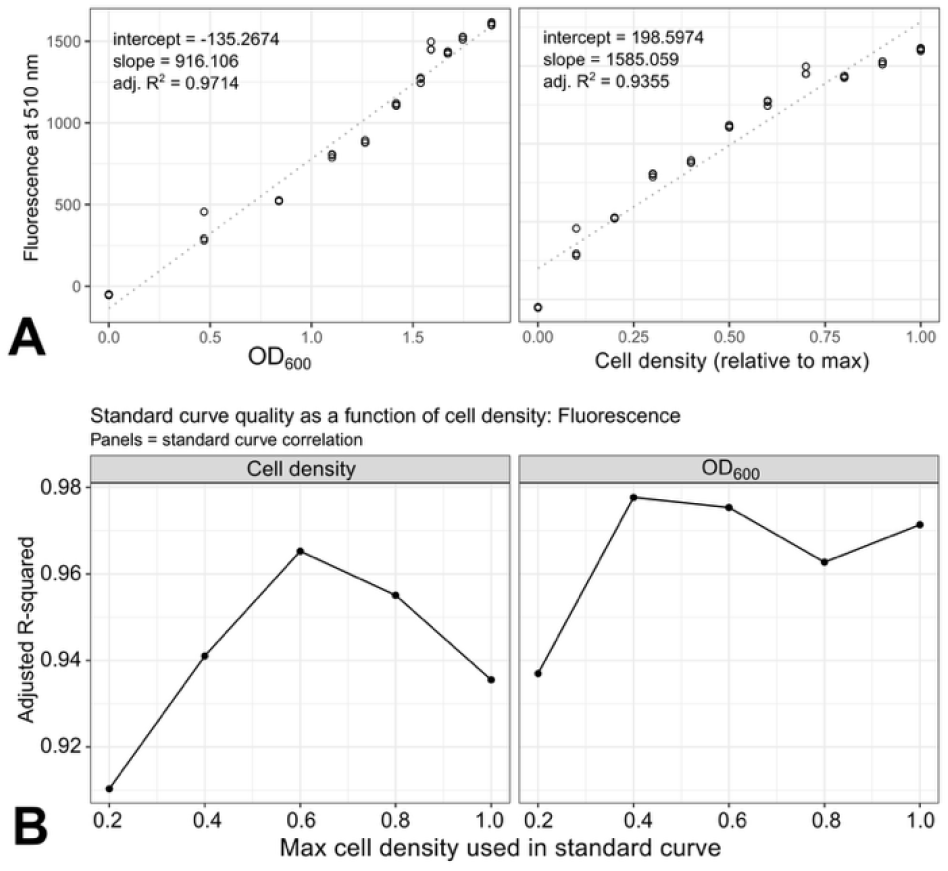
**A)** A strong correlation is observed between fluorescence peak height at 510 nm and OD_600_, suggesting that the former can be used as a proxy for monitoring growth (left). Fluorescence peak height saturates slightly at higher culture densities (right). **B)** Standard curve quality as a function of maximum cell density used in standard curve. Fluorescence shows a better fit to OD_600_ than to cell density.

### 4.4 Spinpod optical system allows measurement of *E. coli* and *S. cerevisiae* growth curves during rotation

As a proof-of-concept demonstration, one growth curve was conducted in a rotating spinpod with *E. coli* and one with *S. cerevisiae (***Fig. 5**). For each growth curve, three full optical spectrum measurements were made (10 seconds per measurement) at each timepoint; one timepoint was taken every 10 minutes (**Fig. 5A–B**). To measure the growth rate of the culture, the height of the peak at 695 nm was extracted from each spectrum measurement (**Fig. 5C–D**), and the corresponding OD_600_ measurement was calculated (**Fig. 5E–F**). This generated growth curves that could be used in a standard microbial growth curve analysis. Fitting to a logistic growth equation, we calculated values of *k* = 9.43 ⋅ 10^−1^, *n*_0_ = 4.84 ⋅ 10^−2^, and *r* = 5.96 ⋅ 10^−1^ for *E. coli* (doubling time: 1.16 hours; 4.89 hours to reach half-max) and *k* = 1.63, *n*_0_ = 1.22 ⋅ 10^−2^, and *r* = 4.44 ⋅ 10^−1^ for *S. cerevisiae* (doubling time: 1.56 hours; 10.1 hours to reach half-max). Note that all values are in units of OD_600_ and hours; the *n*_0_ value is, as expected, extremely low because the initial timepoint of the growth curve was used as the zero value in optical density calculation. The *E. coli* growth curve was not a perfect fit to the logistic curve, showing a few step-changes at approximately 10 hours and 18 hours. These irregularities may be due to biological effects (e.g. clumping in cells) or hardware effects (change in alignment of the optical system).

**Figure 5.**
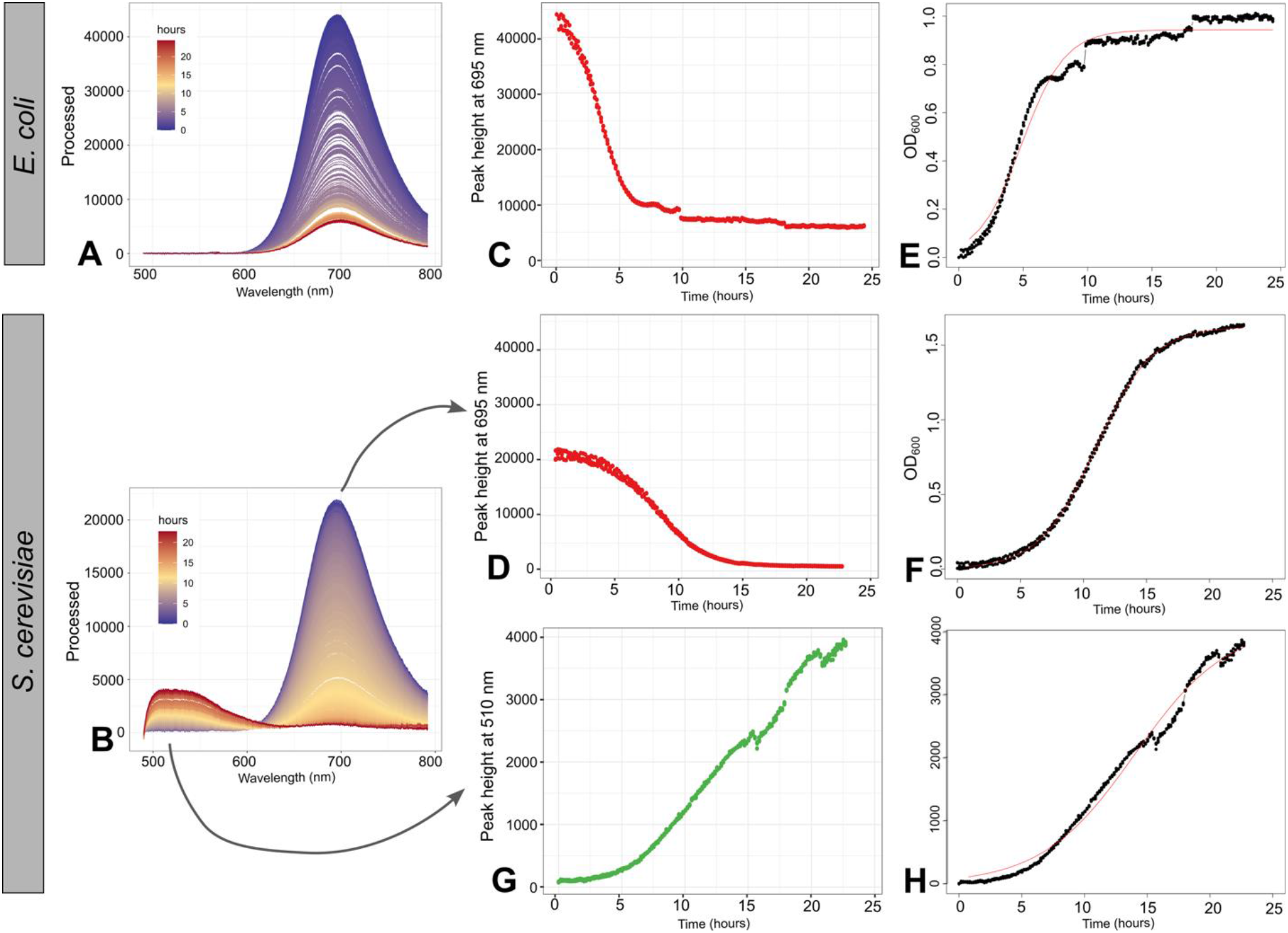
Example real-time optical growth curve data from *E. coli* and *S. cerevisiae* cultures during growth in simulated microgravity. **A–B)** Full optical spectra from 500 nm to 800 nm measured at a frequency of 3 spectra every 10 minutes, for ∼24 hours. Each spectrum is a curve color-coded by the time of acquisition: blue at early timepoints and red at late timepoints. **C–D)** Over time, the height of the peak at 695 nm decreases as increasing cell density decreases the transmission of light from the illuminating LED. **E–F)** Conversion of peak height using a standard curve generates an OD_600_ curve. Black points show the calculated OD_600_ values; red line shows the best-fit logistic curve, generated using the “Growthcurver” package. **G)** In the *S. cerevisiae* curve only, the height of the peak at 510 nm increases over time, as the amount of betaxanthin, which fluoresces green, increases with increasing cell density. **H)** Black points show the peak height over time (identical to panel G); red line shows the best-fit logistic curve.

In addition to optical density, we also demonstrated measurement of betaxanthin fluorescence by *S. cerevisiae* during growth in rotation and fit a logistic growth curve directly to the height of the 510 nm fluorescence peak over time. Fluorescence showed a slower rate of increase than optical density (*r* = 2.700 ⋅ 10^−1^, doubling time: 2.57 hours). Unlike optical density, fluorescence did not appear to plateau during the growth experiment (time to half-max: 14.0 hours), and the maximum fluorescence observed in this experiment was lower than the fluorescence measured for the same density of cells in the standard curve experiment (**Fig. 3**). This could indicate that cells were still producing more betaxanthin at the end of the 24-hour experiment even though cell replication had plateaued. However, more experimentation is needed to fully understand how to interpret fluorescence measurements made in this system. The trend in the 510 nm peak height also showed some anomalies, with step-changes similar to those observed in the *E. coli* OD measurements, around 15, 17.5, and 21 hours.

## 5. Discussion

The spinpod optical system described here represents, to our knowledge, the first system for real-time measurement of growth and activity of microorganisms in simulated microgravity. Taking optical measurements during rotation allows the measurement of fundamental growth parameters such as growth rate and lag time, which are fundamental to microbial physiology experiments, but previously would have required a much more resource-intensive experimental design involving refeeding or destructive sampling.

Here, we demonstrated simple growth curve observations using optical density measurements, and quantification of a constitutively expressed fluorescent pigment. However, the spinpod optical system is versatile: because a whole spectrum is measured at every timepoint, optical absorbance can be measured in any other visible wavelength simply by changing the color of the LED used in the position behind the sample. This would allow measurement of the microbial production of chromogenic compounds (such as carotenoid pigments), or color changes in redox-sensitive or pH-sensitive dyes in the medium that can indicate cellular activity (such as resazurin). The system may also be modified to measure fluorescence of different reporters by changing the excitation light source and the optical filters used, expanding its utility to detecting a wide range of biological activities. Using cells engineered with fluorescent reporters of gene expression, this system could offer a powerful tool for understanding transcriptional dynamics during rotational culture. In a mixed culture of cells carrying different-colored constitutive fluorescent reporters, this system could offer insight into the dynamics of multispecies communities.

Many directions remain open for future development of the spinpod optical system. The current system accommodates only a single experiment at a time; future iterations that allow for multiplexing samples and/or conducting more diverse optical measurements on a single sample would enable more robust and replicated experimentation. Importantly, any new optical measurement made on the spinpod optical system should be benchmarked against measurements made in standard commercial off-the-shelf equipment. Potential confounding optical effects should be considered: for instance, fluorescence measurements are prone to self-quenching, which may be different in a spinpod than in a cuvette or multiwell plate reader.

It is important to acknowledge that microgravity simulators such as spinpods are not complete replacements for spaceflight research, as they only approximate microgravity conditions, and do not fully replicate biological responses to relevant radiation conditions [33]. The interpretation of results from rotational wall vessels experiments continues to be controversial; however, they continue to be the most accessible and commonly-used tool for researchers gathering data in preparation for spaceflight experiments. Improving instrumentation by allowing real-time measurements of biological activity during rotation will help researchers design better experiments to understand the true effect of rotational culture on microorganisms and its fidelity to microgravity as an environmental stressor, to maximize the science return from simulated microgravity experiments and facilitate further microbiological research in support of human space exploration.

## Abbreviations

GFP: Green Fluorescent Protein
HARV: High-Aspect Rotating-Wall Vessel
LED: Light-Emitting Diode
OD_600_: Optical Density (of a sample measured at 600 nm)
PBS: Phosphate-Buffered Saline
RWV: Rotating Wall Vessel
TR: Technical Requirement
USB: Universal Serial Bus
YPD: Yeast Peptone Dextrose
SC: Synthetic Complete

## 6. Acknowledgments

This work was supported by a grant from the NASA Space Biology Program (proposal #20-FG20_2-0015) to J. Lee, with additional support from the Defense Advanced Research Projects Agency B-SURE program (contract #N660012324019) to H. Alper. The views, opinions, and/or findings expressed in this study should not be interpreted as representing the official views or policies of NASA, the Department of Defense, or the U.S. Government. We are grateful to Tim Hammond and Holly Birdsall for technical advice and support during the development of the spinpod optical system.

## S. Supplementary Materials

**Table S1.**
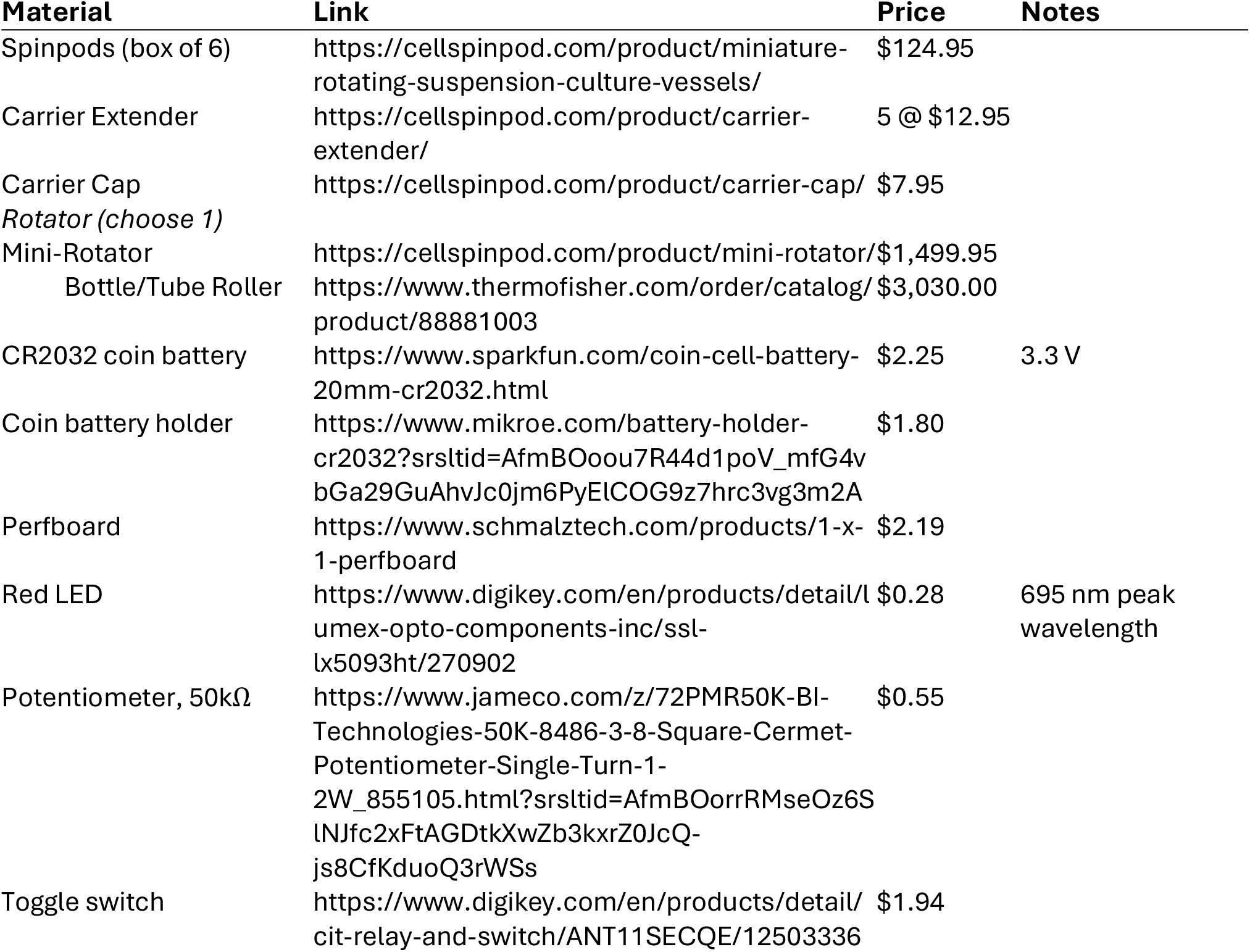
Bill of materials.

